# Vascular Expression of Hemoglobin Alpha in Antarctic Icefish Supports Iron Limitation as Novel Evolutionary Driver

**DOI:** 10.1101/707810

**Authors:** Bruce A. Corliss, Leon J. Delalio, T.C. Stevenson Keller, Alexander S. Keller, Douglas A. Keller, Bruce H. Corliss, Adam C Bjork, Jody M. Beers, Shayn M. Peirce, Brant E. Isakson

## Abstract

Frigid temperatures of the Southern Ocean are known to be an evolutionary driver in Antarctic fish. For example, many fish have reduced red blood cell (RBC) concentration to minimize vascular resistance. Via the oxygen-carrying protein hemoglobin, RBCs contain the vast majority of the body’s iron, which is known to be a limiting nutrient in marine ecosystems. Since lower RBC levels also lead to reduced iron requirements, we hypothesized that low iron availability was an additional evolutionary driver of Antarctic fish speciation. Antarctic Icefish of the family *Channichthyidae* are known to have extreme alteration of iron metabolism due to loss of two iron-binding proteins, hemoglobin and myoglobin, and no RBCs. Loss of hemoglobin is considered a maladaptive trait allowed by relaxation of predator selection, since extreme adaptations are required to compensate for the loss of oxygen-carrying capacity. However, iron dependency minimization may have driven hemoglobin loss instead of a random evolutionary event. Given the variety of functions that hemoglobin serves in the endothelium, we suspected the protein corresponding to the 3’ truncated Hbα fragment (Hbα-3’f) that was not genetically excluded by icefish, may still be expressed as a protein. Using whole mount confocal microscopy, we show that Hbα-3’f is expressed in the vascular endothelium of icefish retina, suggesting this Hbα fragment may still serve an important role in the endothelium. These observations support a novel hypothesis that iron minimization could have influenced icefish speciation with the loss of the iron-binding portion of Hbα in Hbα-3’f, as well as hemoglobin β and myoglobin.

## Introduction

The waters of the Southern Ocean are the coldest on Earth, with temperatures beneath the surface varying between −1.9 and +1.5 Celsius (Sidell and O’Brien 2006). Such frigid conditions are lethal for most mammals, in which blood plasma will quickly freeze (Verde et al. 2007), yet this marine environment hosts many fish species in an ecosystem that exhibits an unexpectedly high amount of biomass despite the harsh conditions (Lam and Bishop 2007; Thresher et al. 2011). Species of the suborder Notothenioid make up 90 percent of the fish biomass in the seas surrounding Antarctica (Sidell and O’Brien 2006), and to withstand cold conditions, they have evolved adaptations such as decreased concentration of red blood cells to minimize blood viscosity, with hematocrit correlating with temperature tolerance across species (Beers and Sidell 2011). An underappreciated consequence of reduced hematocrit is a significant decrease in utilization of elemental iron, since 70 percent of red-blooded mammals’ iron is found in red blood cells in the oxygen-carrying protein hemoglobin (Andrews 2000). Iron is not only considered an essential nutrient for use in various iron-binding proteins (Cairo et al. 2006), but is also established as a limiting nutrient in many aquatic ecosystems (Martin and Fitzwater 1988) including the Southern Ocean (Thomas 2003), where addition of iron is sufficient to induce transient spikes in biomass (Buesseler et al. 2004). Other limiting nutrients have been shown to contribute directly to natural selection in bacteria and yeast (Lewis et al. 1986; Merchant and Helmann 2012). Therefore, we propose that iron limitation could be a selection pressure resulting in adaptations associated with iron binding proteins in the oceans surrounding Antarctica.

Out of all the Notothenioids, Antarctic Icefish of the family *Channichthyidae* are canonically known for the most extreme alterations in iron requirement, lacking expression of hemoglobin, and in half of all species, also myoglobin (Kock 2005a). In this family of icefish, oxygenation is thought to occur purely through diffusion-based transport of dissolved oxygen in the blood (Sidell and O’Brien 2006). The high energetic cost of circulating blood at a rate sufficient for diffusion-based oxygen transport has led previous research to conclude that hemoglobin loss is a net-negative or neutral trait that evolved by chance and remained due to relaxed predator selection (Sidell and O’Brien 2006). However, preservation of such a deleterious trait, even when paired with relaxed selection pressure, is not consistent with the extreme cardiovascular adaptations found in icefish that are required in order to compensate for such inefficient diffusion-based oxygen transport (Kock 2005a). Rather than the random loss of a beneficial trait followed by selection for several necessary compensatory traits, it is plausible that these traits are the product of directional selection resulting from an environmental factor such as limited iron availability.

Knock-out of hemoglobin occurred with the complete genomic deletion of hemoglobin beta (Hbβ), and partial ablation of hemoglobin alpha (Hbα). Given Hbα’s significant role in modulating endothelial nitric oxide signaling (Straub et al. 2012a) in vertebrates, independent of its function in blood oxygen transport, as well as the preservation of a 3’ fragment of hemoglobin alpha (Hbα-3’f) across the genomes of all icefish (Near et al. 2006), we examine whether Hbα-3’f is actively expressed in icefish tissue. While this gene fragment has been classified as an inactive pseudo-gene, we present evidence that Hbα expression in the endothelium has been preserved in icefish retina. This result prompts a reconsideration of whether Antarctic icefish are truly a complete hemoglobin knockout, and reveals the limitation of iron as a possible novel selection pressure in aquatic Antarctic environments. Studying hemoglobin expression in Antarctic icefish may yield insights into how icefish avoid pathological consequences from heightened endothelial nitric oxide production seen in other species and could inform future therapeutics modulating this fundamental vascular signaling pathway.

## Materials and Methods

### Animal Collection & Sample Preparation

Two species of Antarctic notothenioid fishes were collected from the waters of the Antarctic Peninsula region during the austral autumn (April-May) of 2009. *Champsocephalus gunnari*, an icefish species, was caught by otter trawls deployed from the ARSV *Laurence M. Gould* at water depth of 75-150 m in Dallmann Bay (64°08’S, 62°40’W). *Lepidonotothen squamifrons*, a red-blooded notothenioid, was collected in baited traps set at a depth of 200-500 m in both Dallmann Bay and Palmer Basin (64°50’S, 64°04’W). Animals were transferred to the US Antarctic research base Palmer Station, where they were maintained in flowing-seawater aquaria at ambient water temperatures of 0° ± 0.5°C prior to sacrifice. Individuals were first anesthetized in MS-222 in seawater (1:7,500 w/v), and then killed by cervical transection. Retinal tissues were excised quickly, frozen in liquid nitrogen, and stored at −80°C until use. All research was in compliance with the University of Alaska guidelines for work conducted on vertebrate animals (institutional approval 134774-2) and endorsed by the University of Maine (UM) Institutional Animal Care and Use Committee.

### Whole mount & Immunostaining

Retinal samples were thawed from storage at −80°C for 30 minutes or until equilibrium reached with room temperature. Samples were placed in a petri dish, freezing medium drained, and incubated in 4% PFA for 40 minutes, and washed 3 times with PBS for 5 minutes each. Tissue was then flat mounted on a microscope slide outlined with a hydrophobic pen (Sigma-Aldrich Z377821). For staining, samples were blocked and permeabilized with 1 mg/mL Digitonin (Sigma-Aldrich D141) with 10% normal donkey serum (Jackson ImmunoResearch Laboratories 017-000-121) for 3 hours. The following primary antibodies were applied in Digitonin and samples incubated overnight: rabbit anti-hemoglobin beta (1:400, Abcam cat #), rabbit anti-hemoglobin alpha (1:200, Abcam cat 102758), rat anti-CD31 (1:300, Biolegend 102504), and IB4 Lectin conjugated to Alexa Flour 488 (1:200, ThermoFisher 121411). Samples were washed with 0.2% saponin in PBS (Sigma-Aldrich S7900) and incubated overnight with the follow secondary antibodies in 1 mg/mL Digitonin: Donkey anti-rabbit (1:500, Abcam ab150155), Donkey anti-rat, DAPI (1:200, ThermoFisher D1306). Samples were then washed in 0.2% saponin in PBS 6 times for 30 minutes for two days and the mounted with a coverslip, sealed with nail polish.

### Imaging

samples were imaged on a laser point-scanning confocal microscope (Nikon Eclipse TE2000-E Confocal). Z stacks were acquired with a 20x/0.6 oil lens, using a 488 nm laser paired with a 515/30 bandpass, a 546 laser with a 590/50 bandpass, and 647 nm laser with a 650 long pass filter. Fluorophores were excited and imaged sequentially with each laser and filter combination to minimize crosstalk with 1024 pixel resolution and saved as 8-bit images.

## Results

### Vascular Network and Hbα fragment Localization via Wholemount

First we mapped the predicted Hbα-3’f onto the hemoglobin alpha protein (blue; Figure 1) and identified topographically the protein was lacking the heme-binding region. Next, we mapped an alpha globin antibody on the predicted Hbα-3’f (magenta; Figure 1).

**Figure 1:**
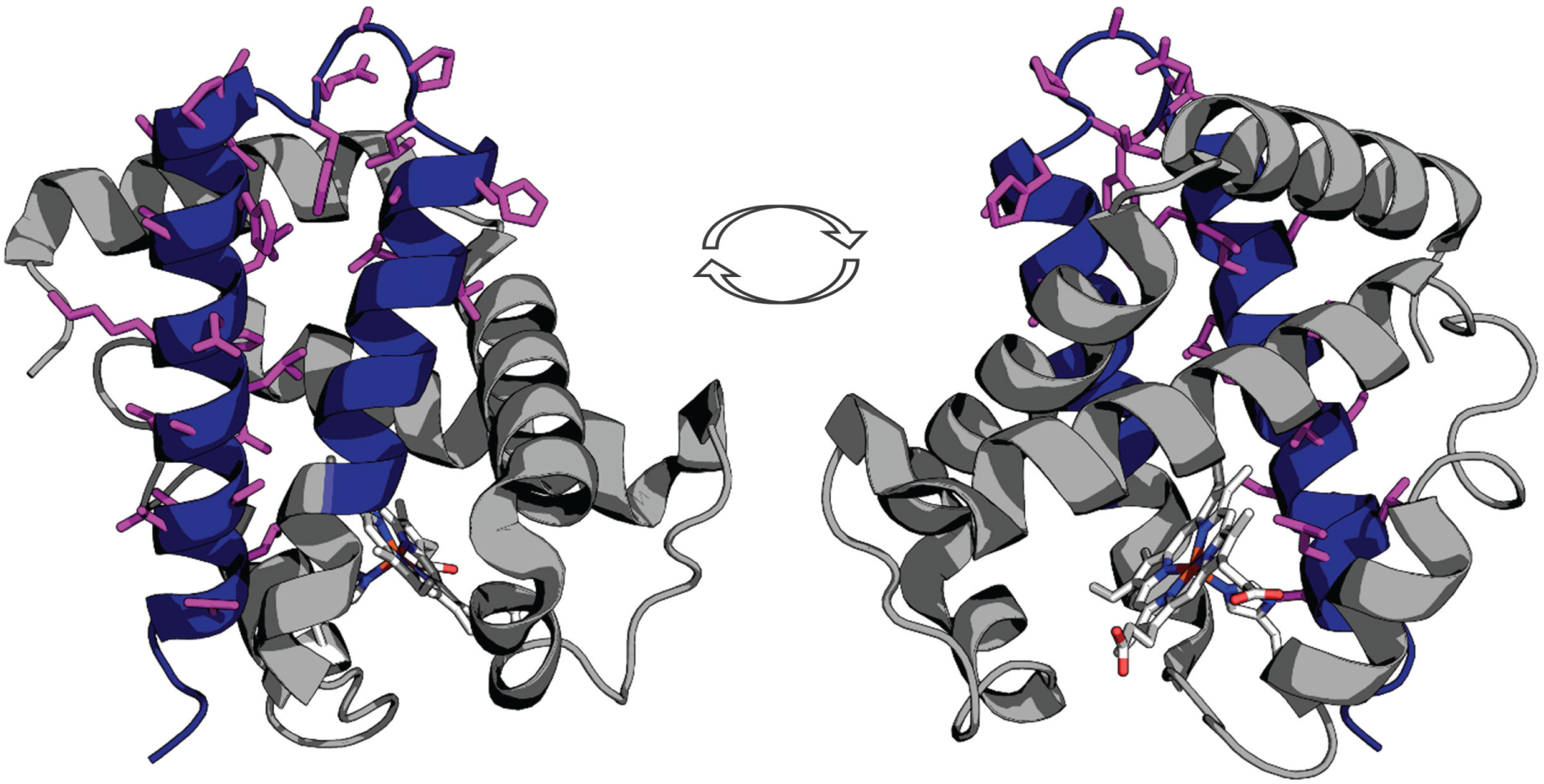
Representation of the truncated icefish alpha globin protein (blue) imposed on human deoxyhemoglobin (gray). Representation of the truncated icefish alpha globin protein (blue) imposed on human deoxyhemoglobin (gray). The epitope for the antibody used in immunofluorescence is represented by magenta side chains, and the entire epitope lies within the truncated gene region. The prosthetic heme group is shown in elemental color scheme. Truncation of the hemoglobin in icefish removes residues critical for heme group stabilization and O_2_ binding ability. The views are a 180° degree rotation of the alpha globin molecule. Using PDBid: 2HHB.

Using our antibody against the Hbα-3’f, we investigated where Hbα protein was localized in whole-mount retinal tissue. In the hyaloid vessels of the vitreoretinal interface of *C. gunnari*, Hbα expression was localized within the vessel wall, denoted by CD31 and IB4 lectin, in blood vessels of all sizes (Figure 2A). As a negative control, there was no detectable expression of Hbβ as expected with its ablation from the icefish genome (Near et al. 2006) (Figure 2B), and no comparable signal observed in unstained tissue with only the secondary antibody present (Figure 2C).

**Figure 2:**
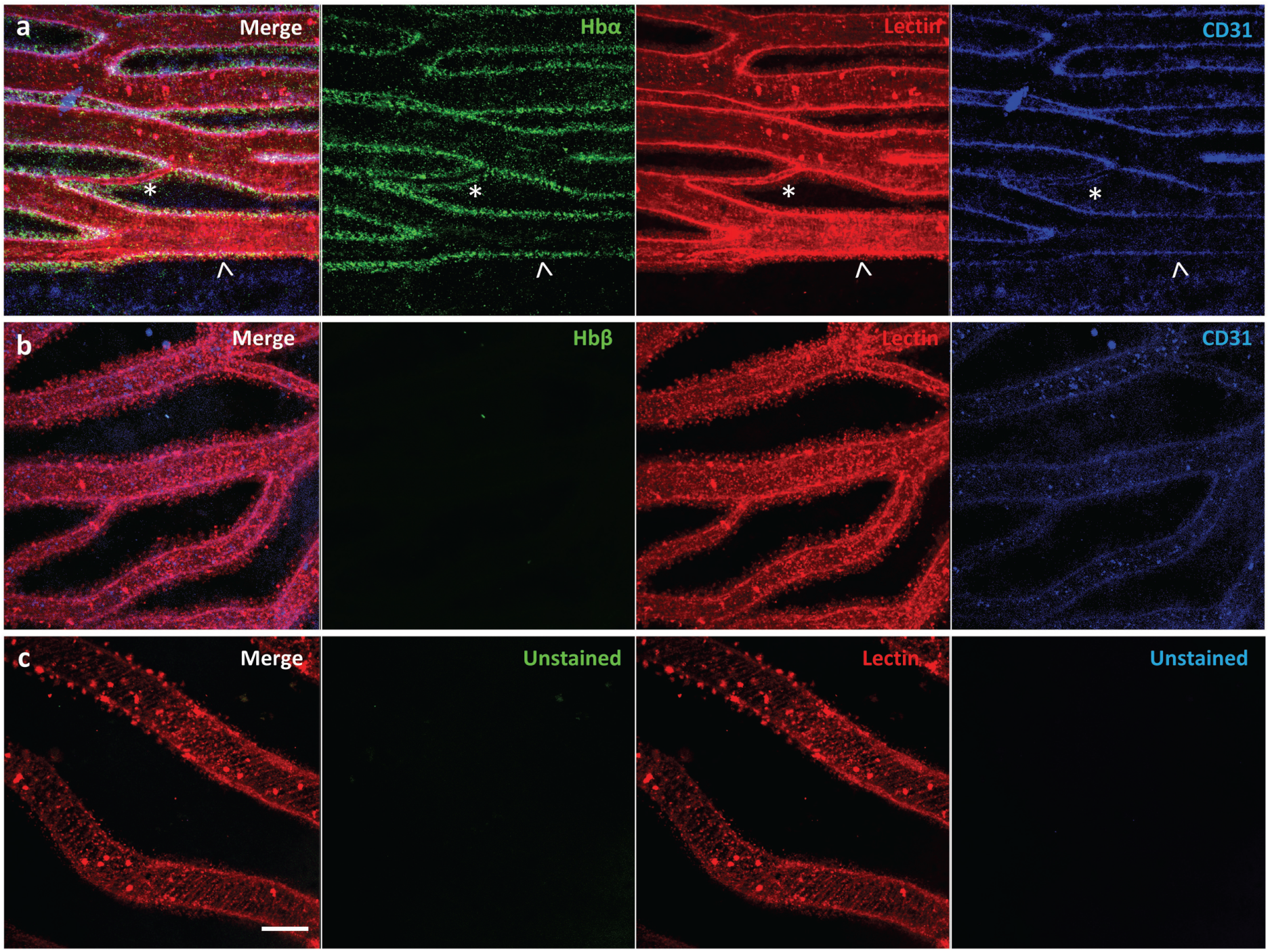
Icefish (*Champsocephalus gunnari)* hyaloid endothelial cells express hemoglobin alpha, while lacking hemoglobin beta expression. (**A**) Retinal surface labeled with anti-Hb! (green), IB4 lectin (red), and anti-CD31 (blue), with a network comprised of both large (arrow) and small (star) vessels. (**B**) Retinal surface labeled with anti-Hb” (green), IB4 lectin (red) and anti-CD31 (blue). (**C**) Retina with lectin as stain control for other channels. Scale bar 100 um, images acquired with 20x/0.75 objective.

Whole mount and immunostaining of *C. gunnari* retina revealed a dense network of IB4 lectin labeled hyaloid blood vessels in the vitroretinal interface radiating from a central optic disk (Figure 3). Of note were the large luminal diameters of the vessel network, with the smallest capillary diameter approximately 30 micrometers and the diameters of the primary vessels ranging from 30 to 150 micrometers.

**Figure 3:**
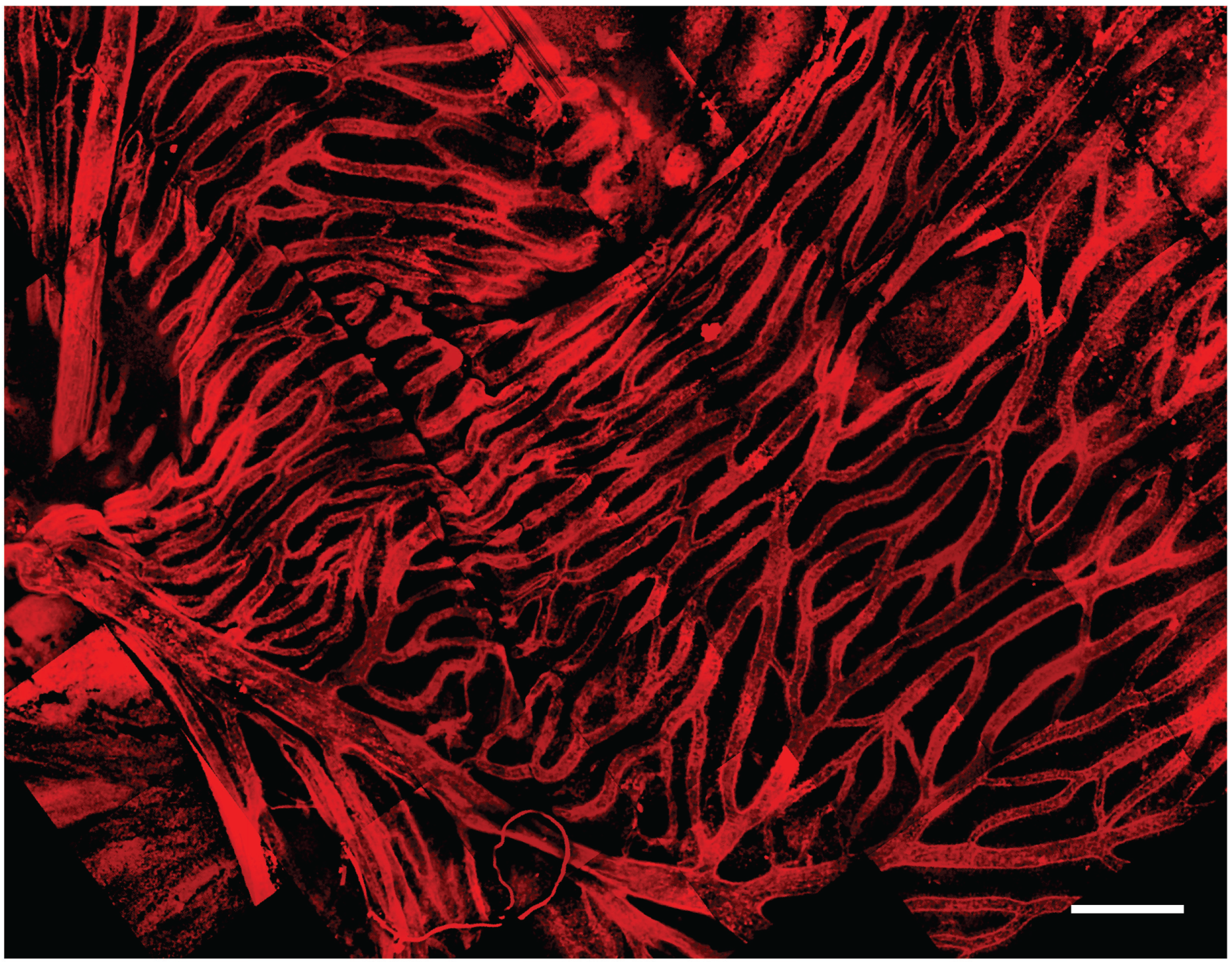
Hyaloid vascular network in vitreoretinal interface of icefish (*Champsocephalus gunnari*). Vasculature labeled with IB4 lectin (red). Scale bar 1 mm, Images acquired with a 20x/0.5 objective.

To confirm that Hbα expression is present in more than one notothenioid, the retina of *Lepidonotothen squamifrons* was also examined. Wholemount immunostaining revealed Hbα expression localized to the endothelial cells contained within the vessel wall of the vasculature residing at the vitroretinal interface (Figure 4A). Similar to *C. gunnari*, there was no detectable Hbβ expression in *L. squamifrons* (Figure 4B), and no comparable signal was observed in unstained tissue with only the secondary antibody present (Figure 4C).

**Figure 4:**
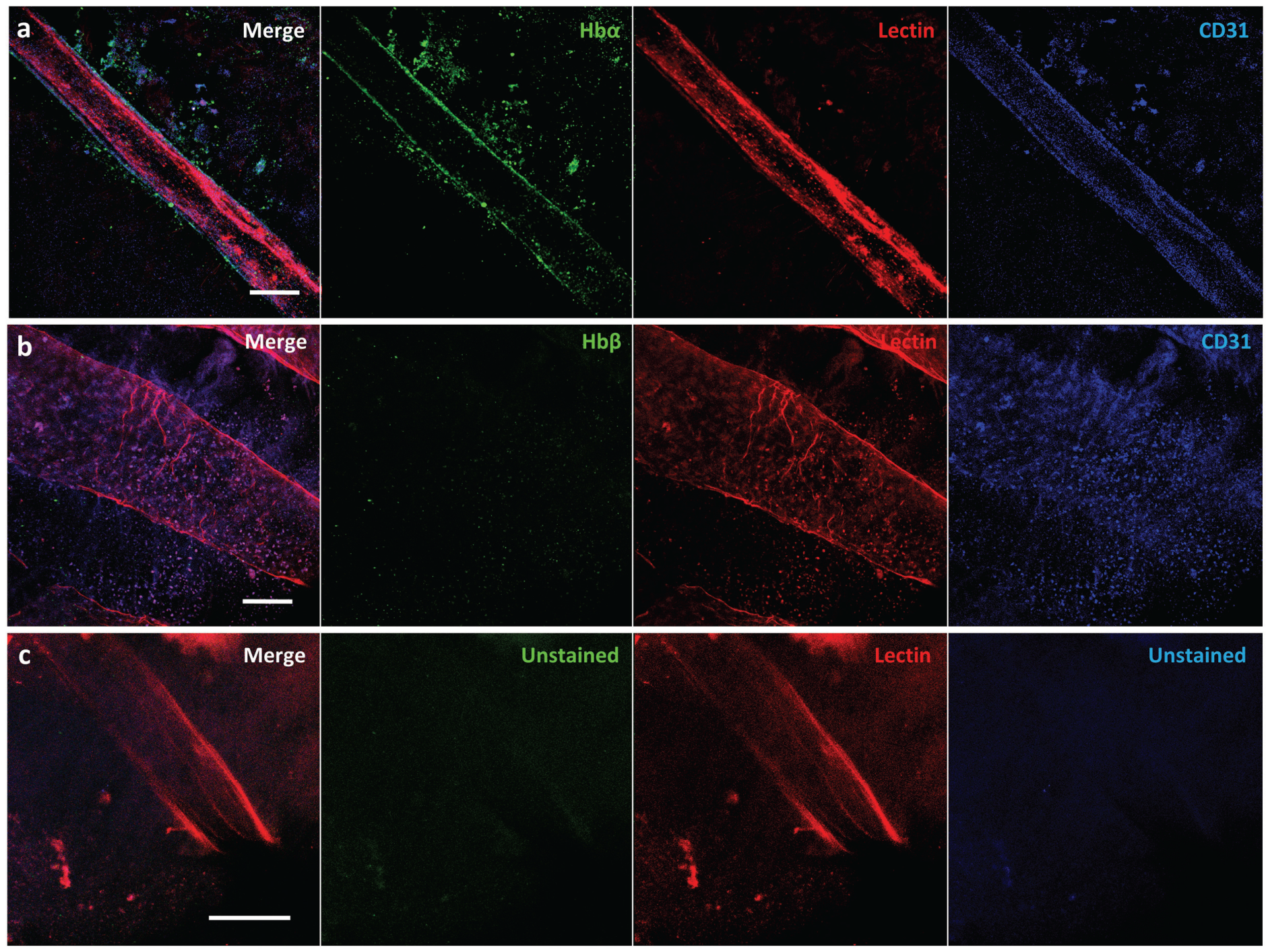
Red-blooded notothenioid (*Lepidonotothen squamifrons*) hyaloid endothelial cells express hemoglobin alpha, while lacking expression in hemoglobin beta. (**A**) Retinal surface labeled with anti-Hb! (green), IB4 lectin (red), and anti-CD31 (blue). (**B**) Retinal surface labeled with anti-Hb” (green), IB4 lectin (red) and anti-CD31 (blue). (**C**) Retina labeled with lectin as stain control for other channels. Scale bar 100 um, images acquired with 20x/0.75 objective.

## Discussion

Our data demonstrate using high-resolution confocal microscopy that fish devoid of RBCs and the genetic deletion of myoglobin and hemoglobin β, express the alpha hemoglobin fragment (Hbα-3’f) in their endothelium. This remarkable discovery demonstrates that alpha hemoglobin localization to endothelium is not confined to mammalian species, and more importantly, it may have more broad implications that originally thought. Some of these concepts are discussed below.

The frigid temperatures of Antarctica have contributed to numerous adaptations for organism to survive in low temperatures (Whittow 1987; Cossins and Macdonald 1989). The Southern Ocean is an especially unique environment because frigid temperatures lead to an oxygen-rich environment with near maximal oxygen saturation (Kock 2005b). This unique combination of extreme environment conditions, paired with relaxation of selection from predation from a historical a lack of apex predators on the food chain (Cocca et al. 1995), has led to especially extensive and unique temperature adaptations compared to more temperate locations.

Adaptations in the fish of the suborder Notothenioid, which make up 90% of the biomass in the Southern Ocean, include the formation of antifreeze glycoproteins that confer resistance to freezing (Coppes Petricorena and Somero 2007) and enzymes optimized for activity at low temperatures to maintain metabolism in frigid conditions (Coppes Petricorena and Somero 2007). These adaptations are mirrored by selection against traits for heat tolerance, such as the loss of functional heat-shock response genes found in other fish species that allow survival at warmer temperatures (Coppes Petricorena and Somero 2007). These highly specialized adaptations to cold come at a cost, however, in that they make notothenioids highly stenothermal, able to survive only in a narrow temperature range from approximately −1.86 to 4°C (Ostadal and Dhalla 2012).

Antarctic icefish of the family Channichthyidae are even more stenothermal than the other members of the suborder Notothenioidei (Cheng and William Detrich 2007; Mueller et al. 2011: 11), with noticeable stress to the organism outside of the −2 to 2 °C range (Sidell and O’Brien 2006). These species are unique in the animal kingdom, exhibiting complete loss of red blood cells and Hbβ, as well as nearly complete loss of Hbα (Sidell and O’Brien 2006). Yet the Channichthyidae family inhabits much of the same aquatic environment as the other notothenioids, typically between 800m and 1,500m depth below sea level (Kock 2005b). Since they coinhabit the same environment, comparing the evolution and physiology of icefish to those of closely related red-blooded notothenioids may yield insight into their diversification. This examination may reveal whether other evolutionary factors, such as minimization of the limiting nutrient iron, played a role in the unique adaptations found these fish and other species of the Southern Ocean.

### Iron Flux Hypothesis: Iron Limitation as a Notothenioid Evolutionary Driver

Similar to the hemoglobin loss found in channichthyids, other notothenioids have evolved a low red blood cell count to counteract the approximately 40% increase of blood viscosity as salt water temperatures near freezing (Near et al. 2006). Hematocrit correlates robustly with thermal tolerance across species (Beers and Sidell 2011). The loss of oxygen carrying capacity that results from reduced hemoglobin expression is viable in cold water environments because oxygen saturation in saltwater nears maximal levels as temperature approaches freezing (Mel’nichenko et al. 2008). Since hemoglobin (with iron bound) makes up 90% of the dry weight content of red blood cells (Rishi and Subramaniam 2017), iron levels in an organism correlate with hematocrit. Hematocrit levels for many of the Notothenioids species are often below 25% (Beers and Sidell 2011), with Antarctic icefish lacking hematocrit entirely, compared to 40-50% typically found in humans.

Iron is critical for several basic biologic functions, including cellular aerobic respiration, oxygen transport through the circulatory system via hemoglobin, and myoglobin function in skeletal muscle (Kaplan and Ward 2013). In humans and red-blooded vertebrates, approximately 70% of the body’s iron content is found in hemoglobin in red blood cells (Andrews 2000), 15% in myoglobin in muscle tissue (Kaplan and Ward 2013), and 6% in other proteins essential for cell metabolism, neurotransmission, and immune system function, with the remaining 9% kept in reserve. An organism with 25% hematocrit would have as much as a 35% reduction in iron requirements. Considering the additional storage and trafficking requirements needed to supply iron for higher hematocrit (Gammella et al. 2014), Notothenioids could have a reduction in iron demand approaching 50% of that needed by many temperate fish species (Gallaugher and Farrell 1998).

In oceanography, the Iron Hypothesis posits that iron is a limiting nutrient in oceanic ecosystems, sufficient to produce phytoplankton blooms on a large scale (Martin et al. 1994). Iron has been demonstrated as a limiting nutrient for biomass in a multitude of open ocean experiments, including the Southern Ocean (Conway et al. 2015). Arctic oceans are especially known to have deficiencies in iron content and flux (Street and Paytan 2005), resulting from the limited input from benthic sediment, atmospheric deposition, and icebergs, alongside limited trafficking of iron between vertical layers of ocean waters (Graham et al. 2015). There have been previous documented cases of limiting nutrients serving as a driver for evolution with plants (Lynch and Brown 2001; López-Bucio et al. 2003; Rennenberg and Schmidt 2010) and microorganisms (Lewis et al. 1986; Merchant and Helmann 2012), providing ample precedent for the possibility that iron limitation may be an underappreciated driver for of Antarctic aquatic species. Mammals have all developed highly specialized iron-binding proteins that act as means of transport and storage (Ganz and Nemeth 2012), offering further demonstration that iron availability can be an significant evolutionary driver. Such extensive biological machinery is necessary because iron is an essential nutrient for all vertebrates (Chen and Paw 2012), and while plentiful on the earth’s surface, is found in low amounts in bioavailable forms (Monsen 1988). Additionally, atomic iron must be kept bound to proteins in a chaperoned state because free iron induces free radical formation that can damage tissue (Emerit et al. 2001).

Further evidence of iron minimization adaptations include Southern Ocean phytoplankton, autotrophs that form the base of the aquatic food chain (D’Alelio et al. 2016), that exhibit unique adaptations that reduce biochemical demand and increase the intracellular flux of bioavailable forms of iron (Strzepek et al. 2011), leading to an 80% reduction in iron requirements compared to temperate oceanic species (Lane et al. 2009). This scarcity of iron availability at the bottom of the food chain means that organisms higher on the food chain only receive a fraction of the iron per mass from phytoplankton compared to other environments, hinting at the Southern Oceans’ unique iron flux. Minimization of iron requirements across the food chain could lead to an ecosystem to support more biomass than otherwise possible. Despite the harsh conditions, there is indeed evidence of higher total biomass than expected in the Southern Ocean (Lam and Bishop 2007; Thresher et al. 2011). Comparing biomass production and iron flux between Antarctic and temperate aquatic environments with ecosystem-level modeling awaits confirmation, but may provide insight to unique iron utilization efficiency between them.

We posit that limited iron availability in aquatic Antarctic environments has led to the selection of traits that conserve its use. In warmer aquatic environments, reducing hemoglobin iron content comes at a steep cost of oxygen-carrying capacity, aerobic respiration ability, and overall organismal fitness. In frigid environments, however, organisms that minimize red blood cell count and iron content would theoretically have the dual benefits of decreased dependence on iron for biomass support paired with an added benefit of lower blood viscosity from reduced hematocrit. Therefore, the oxygen-rich cold waters surrounding Antarctica are uniquely positioned to encourage decreased independence on iron via tenable trade-offs for organismal survival and species fitness. We propose that iron limitation could be a significant driver of icefish evolution, and possibly of portions of the Antarctic ecosystem as a whole.

### Antarctic Icefish as Model of Extreme Iron Metabolism Adaptations

Icefish from the family *Channichthyidae* are known for especially extreme alterations in iron metabolism, making their phylogenic history ideal for examining the evolutionary drivers related to iron minimization. Analysis of iron metabolism in icefish reveals an organism optimized for low iron requirements. Although we present evidence of the expression of a truncated Hbα fragment in icefish tissue, the iron-binding portion of the protein has been ablated along with the entire Hbβ reading frame in all but one icefish species, with *Neopagetopsis ionah* retaining both hemoglobin subunits but thought to form a nonfunctioning complex (Cocca et al. 1995; Near et al. 2006). Loss of hemoglobin and red blood cells leads to 90% decrease in oxygen-carrying capacity (Wujcik et al. 2007) and up to 40% decrease in blood viscosity (Sidell and O’Brien 2006) compared to red-blooded notothenioids. Oxygen transport is therefore purely driven from passive diffusion of surrounding blood vessels into peripheral tissues, dramatically reducing the ability of the circulatory system to deliver sufficient oxygen (F. Garofalo et al. 2009). Not only is the iron demand from hemoglobin absent in icefish, but 6 of the 16 species of Antarctic icefishes have also lost myoglobin expression, an iron binding protein in muscle tissue used for oxygen storage (Sidell and O’Brien 2006). Intriguingly, previous research has concluded that this myoglobin loss was carried out via four independent events during radiation of the species (Sidell and O’Brien 2006), illustrating what could be a strong diversifying selection pressure on icefish myoglobin expression.

Based on iron distribution of red-blooded vertebrates, exclusion of hemoglobin and myoglobin in an organism could lead up to a 90% reduction in iron demands required for homeostasis. Indeed, without iron-binding hemoglobin, iron content in icefish blood plasma is less than 5% of closely related red-blooded species (di Prisco et al. 2002). Yet there is even further evidence of additional iron minimization beyond loss of hemoglobin and myoglobin: concentrations of non-heme iron in Antarctic icefish plasma are one-sixth of that in closely related red-blood species, and are lower by roughly half across various tissues (Kuhn et al. 2016). With a tissue level reduction in iron content, paired with the knockout of two primary iron binding proteins, the iron requirements of icefish normalized to biomass could be greater than 95% compared to other organisms and awaits confirmation.

### Iron Minimization Explains Antarctic Icefish Hemoglobin loss

The loss of hemoglobin is thought to be a non-beneficial evolutionary event paired with a series of compensatory vascular adaptations meant to counteract the loss of oxygen-carrying capacity (Kock 2005a). An energetic analysis of icefish suggests that cardiac function accounts for 22% of resting metabolic demand in icefishes, compared to around 3% with other notothenioids (Sidell and O’Brien 2006). Consequently, hemoglobin loss is perceived as an energetic net negative, requiring far more energy for circulating the high volume of blood plasma required for sufficient oxygen transport than with hemoglobin-mediated oxygen transport (Sidell and O’Brien 2006). Hemoglobin loss is seen as an evolutionary accident, hypothesized to be caused by the presence of a recombination hotspot within the hemoglobin reading frame (Cheng and William Detrich 2007). This predisposition of the disruption of the hemoglobin gene complex (Cocca et al. 1995), paired with a relaxation of selection pressure from predators and oxygen transport from colder temperatures during the speciation of icefish, allowed for the non-beneficial trait to be passed on (Cocca et al. 1995).

We show that a conserved fragment of Hbα is expressed in the vessel walls of the retina of an icefish species, providing evidence that the protein is translationally active. While previous research uniformly references the complete lack of hemoglobin expression in icefish (di Prisco et al. 2002; Kock 2005a; Sidell and O’Brien 2006; Cheng and William Detrich 2007; Mueller et al. 2011), Hbα expression has only been examined in a single species, with mRNA probed indirectly via southern blot with Hbα cDNA fragments from a related red-blooded species (Cocca et al. 1995). Intriguingly, the protein fragment that is detected excludes known interaction and coordination sites, lacking known binding sites for heme (from Leu(F1) to Phe(G5) (Inaba et al. 1998)), eNOS (amino acid sequence LSFPTTKTYF (Keller et al. 2016)), and the α-hemoglobin stabilizing protein that inhibits Hbα precipitation (Feng et al. 2004) (Figure 1). In addition to the endothelial-specific promoter machinery preserved in icefish, BLAST analysis of the Hbα fragment demonstrates high homology with red-blooded vertebrates and humans (Figure 5), and the fragment has been conserved with the species and other vertebrates throughout the phylogenic tree (Figure 6).

**Figure 5:**
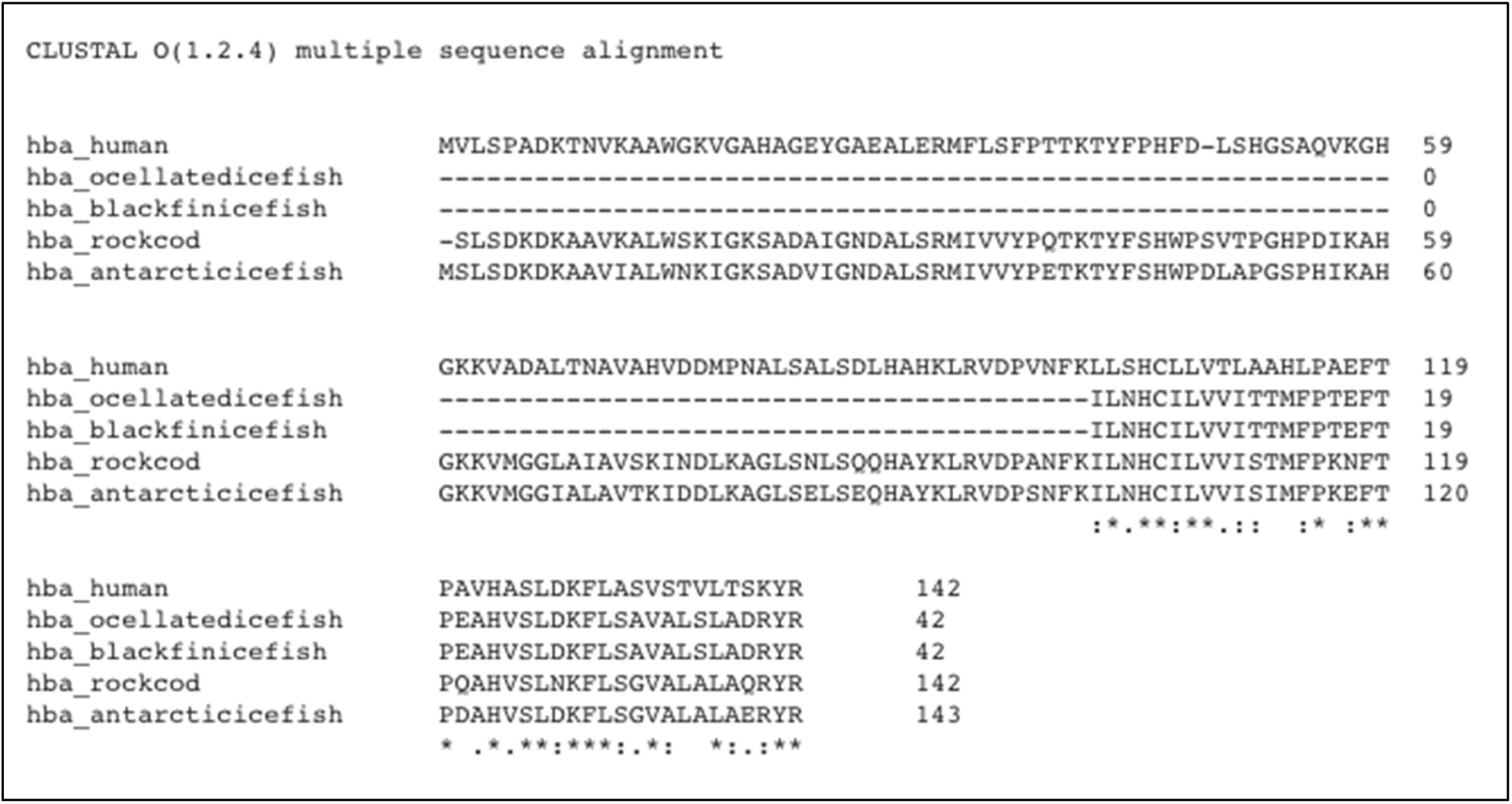
The truncated alpha globin found in icefish is similar to sequences found in other species of fish and the human sequence. Alignment of icefish, icefish-related fish, and human alpha globin sequences using Clustal Omega sequence alignment tool. Symbols *, :, and . represent degree of similarity between amino acids across the sequences. The truncated alpha globin found in icefish is similar to sequences found in other species of fish and the human sequence.

**Figure 6:**
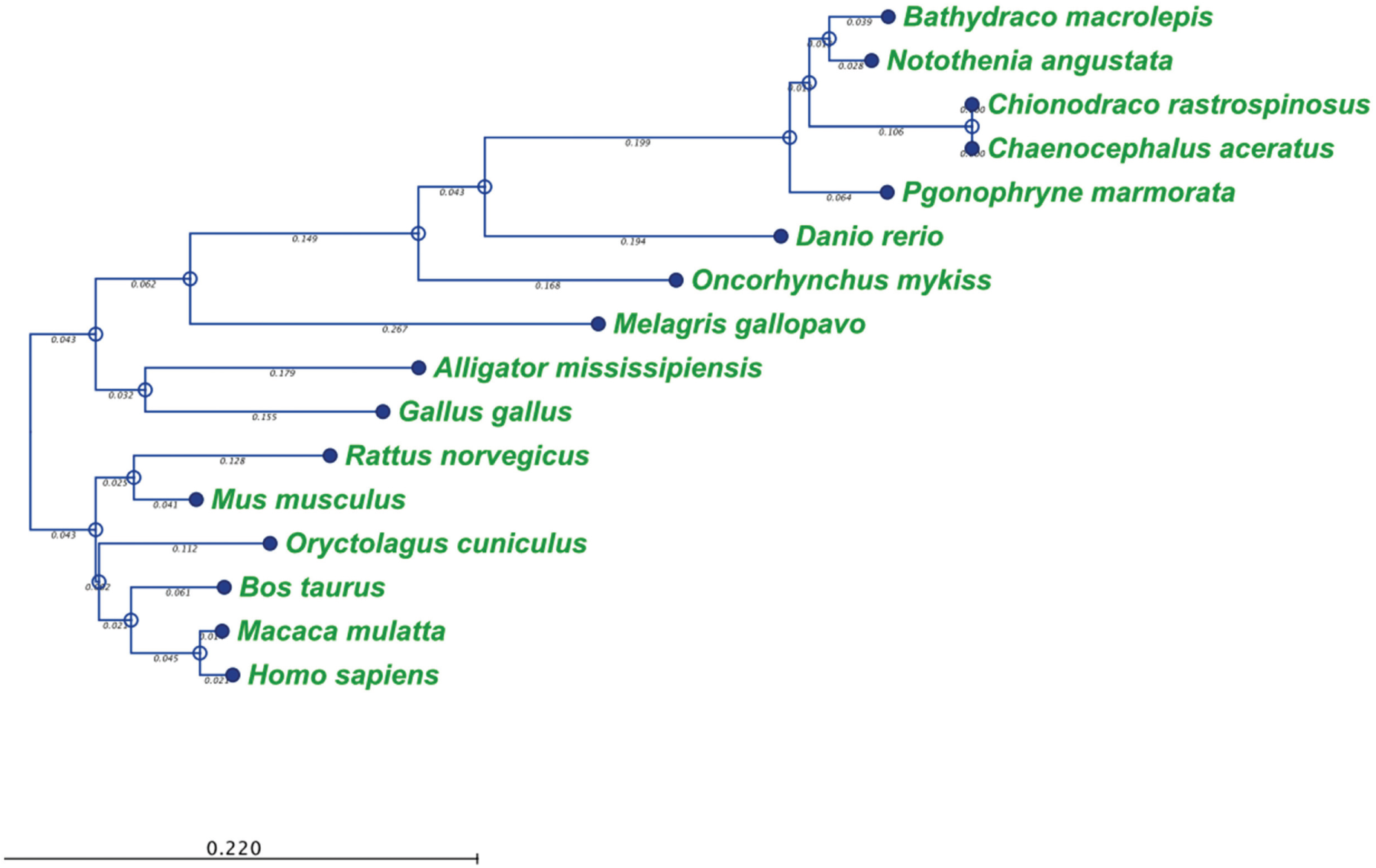
Phylogenic analysis of alpha-globin protein in selected eukaryotic species. Phylogenic tree was constructed using distance-based Neighbor Joining method and protein distance measured with Jukes-Cantor distance adjustment. Scale bar denotes branch length for the amount of change between sequence nodes.

The preservation of functional endothelial-specific expression and conservation of amino acid sequence despite ablation of the majority of the gene suggests that a selection pressure has prevented the complete loss of Hbα. This resistance to complete ablation is most likely explained by Hbα-3’f expression significantly contributing to the fitness of the organism, with complete loss or variation in amino acid sequence being detrimental or even lethal. When considered with the genomic ablation of related iron-binding genes Hbβ and myoglobin, the extreme alterations of three iron binding genes makes it unlikely that these changes are a result of random genetic drift, but implies instead that there was some selection pressure. These changes suggest that rather than a maladaptive (Near et al. 2006) or coincidental neutral benefit trait (Sidell and O’Brien 2006), loss of Hbα could have been the product of diversifying selection pressure, favoring a range of hemoglobin phenotypes (Bargelloni et al. 1998) driven by the recombination hotspot found in close proximity in the genome to Hbα (Cheng and William Detrich 2007), with a niche favoring the near complete loss of the gene.

The existence of an evolutionary driver for hemoglobin loss is further supported by the extreme vascular adaptations required to compensate for the loss of hemoglobin-mediated oxygen carrying capacity. Antarctic icefish exhibit dramatic cardiac hypertrophy (Doake 1987) and a 6-15 fold increase in pump volume compared to other teleosts (Hemmingsen et al. 1972), leading to a dramatic increase in cardiac output (F. Garofalo et al. 2009). Thin, scaleless skin facilitates cutaneous oxygen absorption (Kock 2005a), although its contribution to total oxygen supply is thought to be minor (Doake 1987). Via higher blood vessel density and larger capillary diameter (Wujcik et al. 2007), the icefish vasculature contains 4-fold greater blood volume than red-blooded notothenioids (F. Garofalo et al. 2009), resulting in higher oxygen flux to compensate for the reduced oxygen carrying capacity of diffusion-based oxygen delivery. To minimize oxygen demand, icefish have also evolved lower metabolism (O’Brien et al. 2003; Kock 2005a) and enhanced mitochondrial biogenesis (Coppe et al. 2013).

Building upon prior findings, our results indicate a unique vascular structure of hyaloid vessels at the vitreoretinal interface in icefish. Icefish retinae have been previously visualized on a macroscopic scale through perfusion of opaque silicon rubber and imaged with light photography (Wujcik et al. 2007). Those images revealed a dense hyaloid vascular network branching out from a central optic disk connected to a dense and high-volume capillary network composed of highly isolated branches with few cross-connecting vessels. Higher resolution confocal images of immunostained icefish retina reveal a similar basic vascular network structure, but also a prevalence of smaller vessels connecting vessel branches to form a highly interconnected vessel network. Vessels ranged from 30 µm for the smallest connecting capillaries to 150 µm for the primary vessels emerging from the optic disk. The unusual thickness of these vessels compared to those typically found in vertebrates (Egginton et al. 2002) corroborates previous research demonstrating that mean capillary diameters in Antarctic icefish are 50% larger than capillary diameters in the retina (Wujcik et al. 2007) and skeletal muscles (Egginton et al. 2002) of red-blooded notothenioids. The high density and thickness of the retinal capillary bed is attributed to adaptation to the cold environment of the Antarctic waters (Cheng and William Detrich 2007) to minimize vasculature resistance and maximize oxygen diffusion. The energetic investment needed to maintain such a wide range of adaptations that are required to oxygenate tissues without hemoglobin-mediated oxygen transport further suggests that intense selection pressures are responsible for their initiation and preservation in the gene pool. While there is precedence of complex compensatory adaptations for traits seen as maladaptive, such as the evolution the mammalian retina with sight cells positioned on the far side of the tissue opposite incoming light <u>(Lamb 1995)</u>, there are a lack of cases where a maladaptive trait associated with organism morbidity is retained through a series of compensatory adaptations (Crespi 2000) as seen with icefish hemoglobin loss.

Icefishes co-inhabit the same environments as closely related red-blooded notothenioids, and there is no evidence of any advantages to fitness with hemoglobin loss (Sidell and O’Brien 2006). In fact, evidence points to the reverse, where hemoglobin loss is paired with significant metabolic trade-offs compared to closely related red-blooded notothenioids (Sidell and O’Brien 2006), casting doubt on the possibility of a pure directional selection pressure on hemoglobin concentration, where the extreme phenotype of hemoglobin loss yields a competitive advantage. If the near complete loss of hemoglobin is driven by a diversifying selection pressure, rather than a maladaptive event or neutral drift, then an additional driver could be required to more fully explain why a diversity of phenotypes of hemoglobin concentration are found in fish of the Southern Ocean (Beers and Sidell 2011). We propose that iron limitation may be the missing evolutionary force behind icefish adaptation, and an underappreciated driver with the evolution of many of Antarctic aquatic species in general. We present evidence that supports the notion that a diversifying selection pressure may have driven icefish evolution with hemoglobin-mediated oxygen transport, where frigid temperatures and minimization of blood plasma viscosity are insufficient to explain the driving forces behind icefish evolution. Minimization of iron requirements could contribute to organism fitness when prey populations are restricted or contain reduced iron content, as found with phytoplankton (Strzepek et al. 2011) and fish (Beers and Sidell 2011) in the Southern Ocean. Low iron usage may have aided survival during the crash in Antarctic biodiversity that co-occurred with icefish speciation (Eastman 1993) roughly 8.5 million years ago (Near 2004). The evolutionary importance of conservation of metabolic inputs has precedence that includes adaptations with hibernation of mammals in winter (Geiser 2013), dormant states in bacteria during environmental stress (Watson et al. 1998), and starvation responses found across mammals (Wang et al. 2006).

### Antarctic Icefish as a Model of Heightened Endothelial NO Bioavailability

Recently, hemoglobin alpha, canonically known for its role in oxygen transport via binding with hemoglobin beta in red blood cells, has been shown to modulate vascular remodeling in protrusions of endothelial cells called myoendothelial junctions (MEJ) (Straub, Zeigler, et al. 2014). These regions are on the basolateral membranes of endothelial cells, proximal to smooth muscle cells, and facilitate communication between the two cell types in the vascular wall. Hbα modulates endothelial NO flux at the MEJ (Straub et al. 2012b) in resistance arteries by binding to eNOS and acting as a scavenger of NO (Butcher et al. 2014). Due to NO’s short biological half-life (Thomas et al. 2001), Hbα serves as a significant negative regulator for the availability of endothelial NO reaching proximal smooth muscle cells. Disruption of the Hbα-eNOS interaction can lead to smooth muscle vasodilation and reduction in blood pressure (Keller et al. 2016), while eNOS inhibition leads to vasoconstriction of the peripheral vasculature and can induce significant increases in blood pressure (Li and Förstermann 2000). Indeed, pharmacologically increasing the bioavailability of endothelial NO (Kurowska 2002) is perceived as a promising therapeutic strategy in atherosclerosis (Barbato and Tzeng 2004), ischemia (Barbato and Tzeng 2004), diabetes (Masha et al. 2011), and hypertension (Hermann et al. 2006). Paradoxically, completely unregulated hyperactive endothelial NO generation can be pathological. A pronounced example is a recent preclinical study in rhesus monkeys, where an antibody exhibiting off-target effects leading to elevated NO production (Pai et al. 2016) caused severe systemic vasodilation, as well as hypotension, hematemesis, hematochezia, and morbidity. Additionally, elevated levels of NO is used a biomarker is various diseases (Arkenau et al. 2002; Pham et al. 2003), results in apoptosis (Blaise et al. 2005), produces cytotoxic oxygen radicals, exerts cytotoxic and antiplatelet effects (Sim 2010), inhibits enzyme function, promotes DNA damage, and activates inflammatory processes (Hollenberg and Cinel 2009).

In the absence of Hbα scavenging NO, as evident via the exclusion of the binding sites for NOS and heme in Hbα-3’f, production of NO might be up regulated. Icefish could potentially serve as a model organism to study up regulation of endothelial nitric oxide signaling (Beers and Jayasundara 2015) while avoiding the pathological ramifications that are experienced in Hbα-expressing vertebrates (Pai et al. 2016). A possible function of Hbα-3’f could include NO binding at Cysteine 5 in a similar fashion to the established Cysteine-NO interaction found at Cysteine 93 in Hbβ (Sampath et al. 1994; Helms and Kim-Shapiro 2013). Instead of trapping NO in the Hbα-eNOS complex at the point of generation, NO trapping would be carried on in a diffuse form with the freely disassociated Hbα fragment throughout the cytosol. Altered nitric oxide kinetics could represent a safe method to up regulate nitric oxide metabolites in the vessel wall while still maintaining negative regulation that avoids NO toxicity.

The vascular evolutionary adaptations compensating for loss of heme-mediated oxygen-carrying capacity are thought to be facilitated through nitric oxide signaling (Cheng and William Detrich 2007). Enriched endothelial NO has been shown to play a role in modulation of vasodilation (Palmer et al. 1987), angiogenesis (Ziche and Morbidelli 2000), cardiac hypertrophy (Wollert and Drexler 2002), mitochondria size (Urschel and O’Brien 2008), and mitochondrial biogenesis (Nisoli and Carruba 2006; O’Brien and Mueller 2010), all of which are exaggerated phenotypes found in hemoglobin-lacking icefish (Kock 2005a). Several studies provide evidence of the presence of a functional NOS signaling system (Pellegrino et al. 2004) and expression of eNOS has been preserved in endothelial cells of icefishes (Filippo Garofalo et al. 2009), along with a 50% greater plasma load of NO metabolites (NO_x_) in icefish compared to red-blooded notothenioids (Beers et al. 2010). However, it is important to note that a significant portion of this elevated NO metabolite load could be from a physiologic response to hemoglobin loss, revealed that, at least in a transient fashion, when red-blooded notothenioids were subject to chemically induced anemia that resulted in a 70-90% reduction in hemoglobin concentration, NOx metabolites also increased by 30% (Borley et al. 2010).

Nitric oxide metabolite buildup in icefish is theorized to be caused by reduced degradation rather than increased generation. Previous studies have shown that vascularized icefish tissue has a 50% decrease of NOS (Beers et al. 2010), the primary source of endothelial NO generation, compared to closely related red-blood species. The alteration of Hbα’s heme-based NO scavenging ability in the Hbα −3’f could explain how icefish simultaneously express less NOS but exhibit greater NO load in the vasculature (Beers et al. 2010).

## Conclusion

We demonstrate that Hbα-3’f is expressed transcriptionally, translationally, and localized to the vasculature. Conservation of the Hbα-3’f amino acid sequence between icefish species and red-blooded vertebrates, along with preservation of endothelial-specific promoter machinery alongside loss of all known Hbα interaction regions, suggests that this Hbα-3’f fragment plays a novel, unknown role in the endothelium. These findings demonstrate that icefish do not technically have both hemoglobin genes knocked out, but do suggest that all known Hbα functions have been disabled, including known interaction regions with eNOS, heme, and NO. The ablation of the majority of the Hbα gene may essentially represent a natural mutagenesis experiment where nonlethal portions of the gene are eliminated, offering a possible opportunity to identify a novel role of the Hbα fragment region that may translate back to red-blooded vertebrates.

Preservation of the Hbα-3’f protein fragment suggests that a diversifying selection pressure could have driven the process. We propose that iron is a novel evolutionary driver for icefish hemoglobin loss, and perhaps even for the decreased hemoglobin concentration found in various other Antarctic aquatic species. Testing this hypothesis will require an examination of iron flux and iron utilization on an organism and ecosystem level.

